# Orthogonal dietary niche enables reversible engraftment of a gut bacterial commensal

**DOI:** 10.1101/275370

**Authors:** Sean M. Kearney, Sean M. Gibbons, SE Erdman, EJ Alm

## Abstract

Interest in manipulating the gut microbiota to treat disease has led to a need for understanding how organisms can establish themselves when introduced into a host with an intact microbial community. While probiotic or prebiotic approaches typically lead to a transient pulse in an organism’s abundance, persistent establishment of an introduced species may require alternative strategies. Here, we introduce the concept of orthogonal niche engineering in the gut, where we include a resource typically absent from the diet, seaweed, to establish a customized niche for an introduced organism. We show that in the short term, co-introduction of this resource at 1% in the diet along with an organism with exclusive access to this resource, *B. plebeius* DSM 17135, enables it to colonize at a median abundance of 1%, frequently increasing in abundance to 10 or more percent. We construct a mathematical model of the system to infer that *B. plebeius* competitively acquires endogenous resources. We provide evidence that it competes with native commensals to achieve its observed abundance. We observe a diet-dependent loss in seaweed responsiveness of *B. plebeius* in the long term and show the potential for IgA-mediated control of putative invaders by the immune system. These results point to the potential for diet-based intervention as a means to introduce target organisms, but also indicate potential modes for failure of this strategy in the long term.

## INTRODUCTION

Introducing new bacteria into an intact or disturbed microbial community is one of the primary goals of microbial therapeutics. However, we have limited knowledge about the features that govern successful colonization by introduced microorganisms. Recent work has demonstrated that the presence of functional metabolic capacity beyond that of the endogenous microbiota contributed to persistence of colonization by an introduced *Bifidobacterium* species (Maldonado-Gómez et al., 2016). A strong predictor for engraftment of microorganisms introduced by fecal microbiota transplant is the presence of shared organisms in the donor and recipient (Li et al., 2016; Smillie et al., 2018), likely indicating that shared functional capacity mediate colonization in this case. Further, transposon sequencing of a model gut community of *Bacteroides* species identified arabinoxylan as capable of modulating the abundance of a single strain (Wu et al., 2015). These findings led us to believe that we could enhance the colonization of an introduced organism into an intact ecosystem by providing it exclusive access to a resource unshared by the rest of the community.

We identified a resource unlikely to be used by microorganisms in the lab-mouse gut: the red algae, *Porphyra*, comprising the edible seaweed nori. Nori contains complex, sulfated polysaccharides including porphyran, and unlike terrestrial plants, includes vitamin B12 (Hehemann et al., 2012), which has previously been implicated as a fitness determinant for *Bacteroides fragilis* in the murine GI tract (Goodman et al., 2009). The pathways for degradation of porphyran are rare in the human gut microbiota, and almost exclusively found in metagenomic samples from individuals in populations known to eat seaweed, primarily Japanese individuals, who consume on average 5-10 g of seaweed daily (Hehemann et al., 2012). Horizontal gene transfer of the pathways necessary for breakdown of porphyran from a marine bacterium to *B. plebeius*, a gastrointestinal commensal, suggest strong selective pressures for the acquisition of this trait (Hehemann et al., 2010). Because of these observations, we inferred that expanding the diet of mice to include seaweed would result in a clear signal through increasing the abundance of a narrow range of organisms capable of using this resource. In the absence of a signal, we inferred that the mouse gut microbiota would be open to invasion by *B. plebeius* in the presence of seaweed given its unique access to this resource.

Here, we show that *B. plebeius* colonizes mouse guts at high levels in the presence of a preferred substrate (i.e. polysaccharides in seaweed), with no evidence of competition for this substrate from native gut bacteria. We conduct parametric analysis of a nonlinear dynamical model of the system to explain how low levels of an exclusive resource can lead to abundant colonization of an introduced species. Enhanced colonization, however, comes at a cost to *B. plebeius*, with its levels depleted in the long term depending on the original seaweed treatment. We provide preliminary evidence for microbial competition or host-immune inhibition of this introduced organism in mediating these outcomes. These results provide a first proof-of-principle for orthogonal niche engineering as a method for synbiotic design (i.e. controlling abundance of introduced microorganisms through manipulating resource availability) (Krumbeck et al., 2015; Panigrahi et al., 2017), but also reveal limitations to this strategy. In the context of microbial therapeutics, we show stable engraftment of a non-indigenous bacterial strain into an intact gut community in the presence of its engineered niche. However, our results also indicated that foreign microbes, when persistently maintained at high abundance, might be at a disadvantage when entering a system unaccustomed to their presence due to negative feedbacks from the host immune system.

## RESULTS

*Seaweed treatment does not change the composition of the mouse microbiota* Orthogonal niche engineering requires that introduction of the niche (in this case through provisioning of seaweed), does not favor the growth of organisms already in a community. To validate that organisms in the mouse gut do not have the capacity to outgrow significantly on seaweed-derived substrates, we introduced seaweed into mouse chow and followed the changes in the composition of the microbiota over time. We singly housed six-week old female C57BL/6 mice and randomly assigned them to two groups: (1) those receiving standard mouse chow (control) and (2) those receiving seaweed at 1% in their mouse chow (seaweed) (Fig 1A). Mice received seaweed chow continuously for 16 days, following a 32 day washout period and resumption of seaweed feeding for an additional 8 days to assess within-mouse reproducibility of the effects of seaweed feeding. Fecal samples were collected daily for the duration of the experiment, and temporally separated subsets were selected for V4 16S rDNA amplicon sequencing.

**Figure 1.**
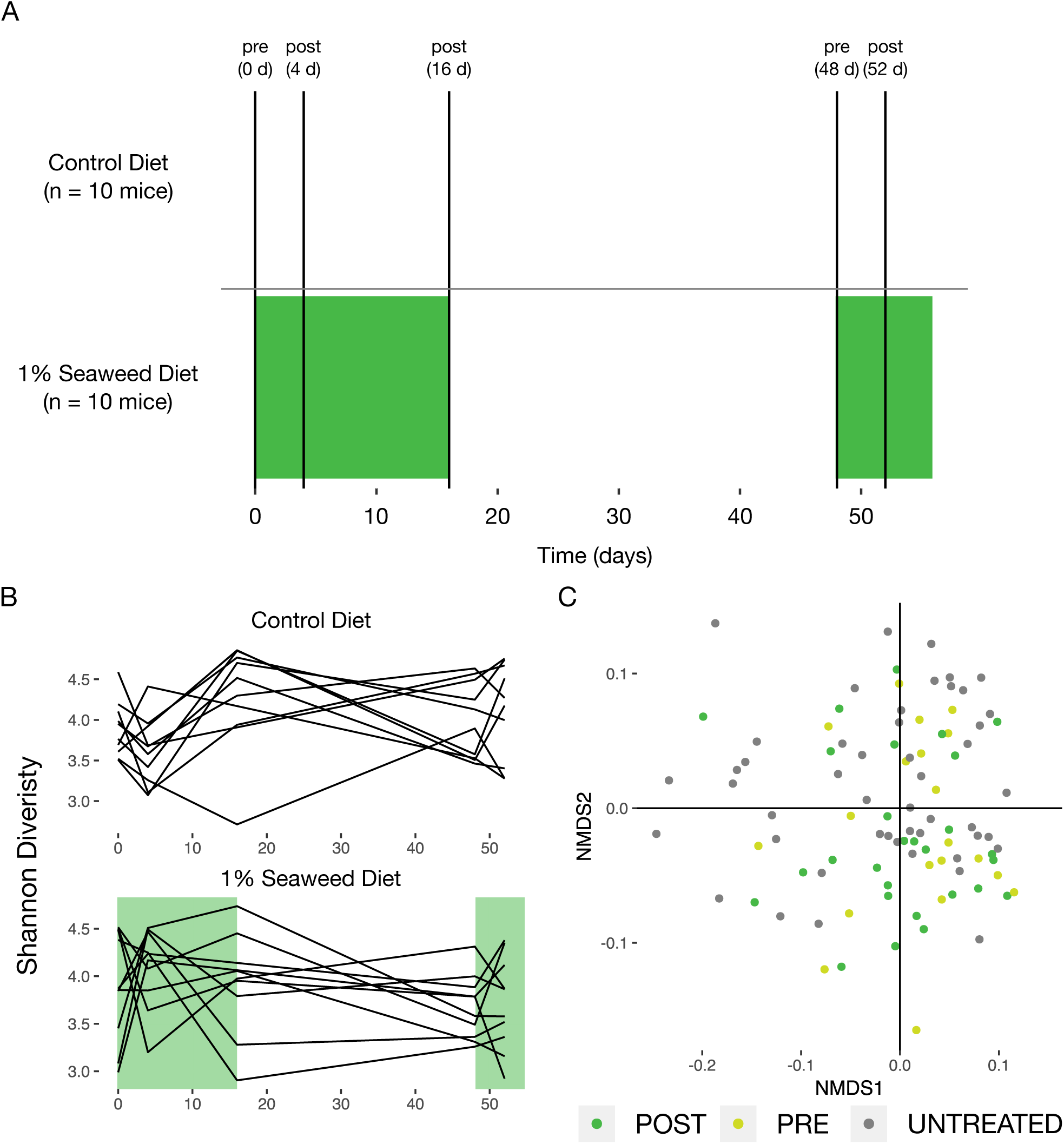
Seaweed does not alter the microbiota of conventional mice. (A) Experimental design: samples were collected at the indicated time points for 16S rDNA sequencing. Green shaded region indicates period of seaweed feeding. (B) Shannon diversity over time for control animals and seaweed treated animals, with green shading again indicating the period of seaweed feeding. (C) Nonmetric multidimensional scaling of the Jensen-Shannon Divergence across mice treated and untreated with seaweed and in pre- and post-time points.

We expected that seaweed treatment would not lead to compositional changes in the microbiota. We examined the change in community structure on a seaweed diet by tracking the alpha diversity over time (Figure 1B). There were no consistent changes in Shannon diversity in the seaweed-treated mice in either the initial seaweed or late seaweed treatment time points, suggesting that seaweed feeding did not coherently alter the mouse gut microbiota in a way reflected in alpha diversity. We used Jensen-Shannon Divergence and non-metric multidimensional scaling to identify whether communities became more similar after seaweed treatment. Seaweed treated communities did not cluster separately from controls, also suggesting that this treatment led to no changes observable in the community at this level (Figure 1C). Additionally, we found no evidence for the presence of genes involved in porphyran breakdown based on qPCR targeting the β-porphyranase gene present in the polysaccharide utilization locus (PUL) of *B. plebeius* (Hehemann et al., 2012), indicating that the genetic potential to use polysaccharides present in the seaweed was absent.

Having shown that the native mouse microbiota has limited responsiveness to seaweed, we reasoned that an organism previously reported to exploit this niche would grow in its presence in the mouse gut. Further, we wanted to pick a strain that would not engraft unless given a fitness advantage. In FMT, most strains do not engraft even though they are directly transferred between human hosts (Smillie et al., 2018), so we expected using a species not native to the mouse gut would satisfy this criterion. In particular, the human-derived *B. plebeius* DSM 17135 strain carries the genes for porphyran degradation, as do at least two other (GI commensal) isolates. This pathway is carried on an integrative conjugative element, and has historically been transferred between gut microorganisms (see, in addition to *B. plebeius, Bacteroides sartorii* JCM 16497 and *Porphyromonas bennonis* JCM 16335). The positive selection on these genes, indicated by their transmission via HGT, suggests that organisms carrying these genes will be more fit in an environment rich in this substrate.

### Introduction of *B. plebeius* into the digestive tract of mice

To assess the feasibility of introducing *B. plebeius* into the GI tract of mice, we ran an experiment with four groups of outbred female Swiss mice co-housed by treatment. The treatment groups were as follows: (1) mice that were gavaged at the beginning of the experiment with 10^7^ CFU of *B. plebeius* (*B. plebeius* only), and mice that were gavaged with *B. plebeius* and (2) fed seaweed continuously (seaweed) or (3) delivered pulses of seaweed with four days on and four days off (pulsed), and (4) mice that were fed seaweed but not gavaged with *B. plebeius* (control) (Figure 2A). We chose outbred mice in order to reduce the likelihood of observing genotype-specific signals in colonization by *B. plebeius*. Animals were co-housed by treatment to improve exchange of *B. plebeius* via coprophagy, to reduce probability of stochastic extinctions, and to select for the fittest genotype across all animals, rather than a single-animal optimized genotype.

**Figure 2.**
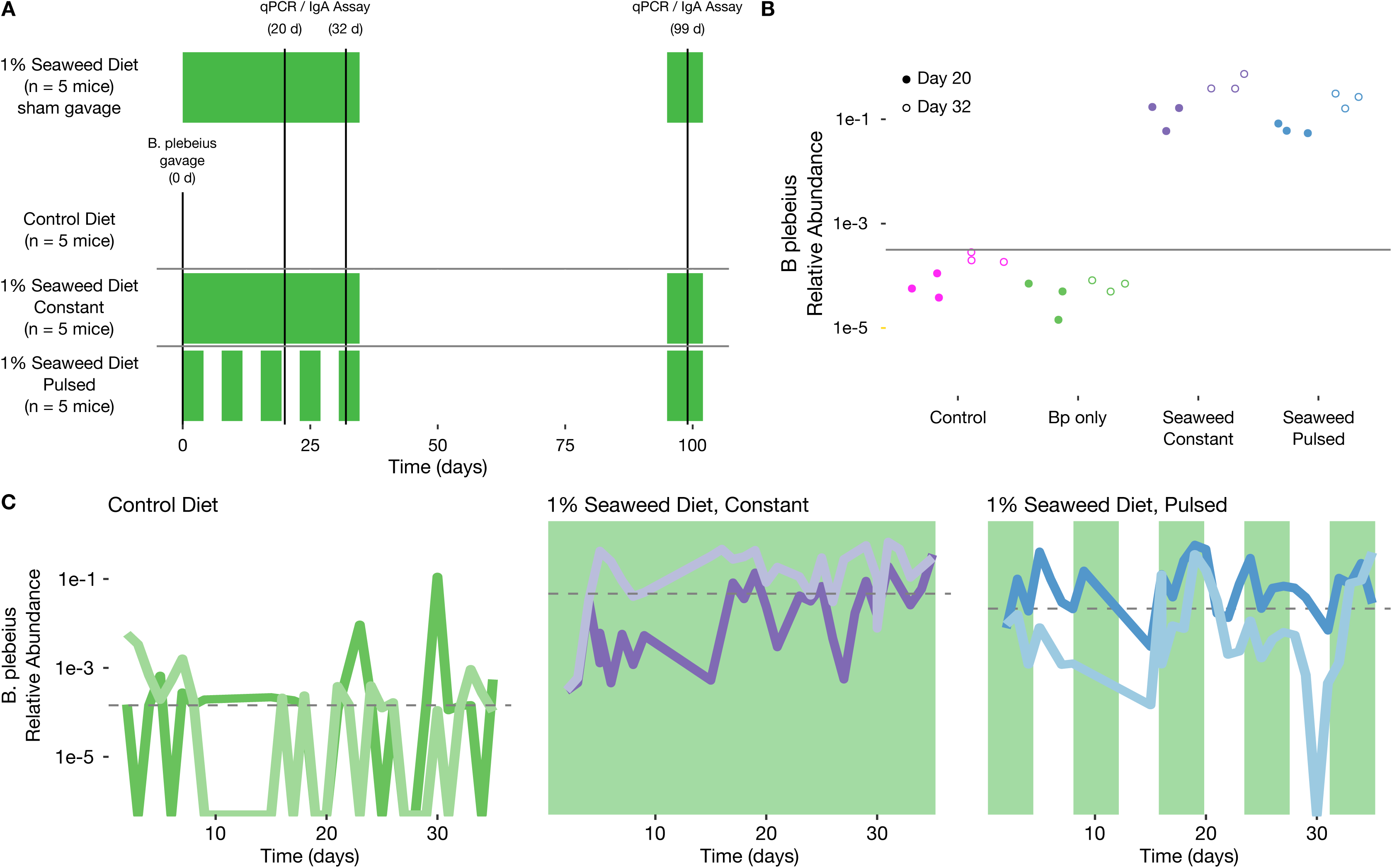
*B. plebeius* reaches persistent high level abundance in mice consuming seaweed. (A) Experimental design: samples were collected where indicated for qPCR and IgA-Sequencing; two animals per *B. plebeius* only, constant seaweed, and pulsed seaweed were sampled densely across until cessation of the first seaweed treatment window. Green shaded region indicates the seaweed dosing windows. (B) qPCR-based relative abundance of *B. plebeius* in control, *B. plebeius* only, constant seaweed, and pulsed groups at indicated time points. The lower limit of detection from non-specific amplification is marked with a gray line. (C) Time series of *B. plebeius* in *B. plebeius* only, constant seaweed, and pulsed groups, line colors distinguish individual animals, green shading indicates seaweed-dosing periods.

From each mouse, fecal samples were collected daily for a 35-day period, and a subset of these samples was processed for V4 16S rDNA amplicon sequencing. Follow-up samples from 2 months after cessation of the initial seaweed feeding experiment were collected to determine whether *B. plebeius* persisted in mice feces long after the initial treatment. During the experiment we did not note any adverse affects on the mice across the treatment groups.

### *B. plebeius* abundantly colonizes the GI tract of mice feeding *ad libitum* on seaweed

We focused our sequencing efforts on the time series of 2 mice per treatment group, each randomly selected from the gavaged and untreated (*B. plebeius* only), gavaged and treated with 1% seaweed (seaweed), and gavaged and treated with pulsed 1% seaweed (pulsed) groups. In the seaweed treated groups, we expected to find enrichment for *B. plebeius* and dilution and eventually extinction in the absence of seaweed. In the pulsed treatment, we expected that *B. plebeius* levels would rise and fall to track the presence of the seaweed.

In fact, we found that the relative abundance of *B. plebeius* increased by more than two orders of magnitude (p = 0.001, n = 10 (seaweed + pulsed) mice, n = 5 *B. plebeius* only mice, Mann Whitney U test) in the seaweed and pulsed groups relative to the *B. plebeius* only group, suggesting that seaweed treatment enriches for *B. plebeius* in the GI tract of these mice (Figure 2B and Figure 2C). Indeed, at some points, *B. plebeius* rises in abundance to nearly 50% of the whole community. As expected, we observe drops (albeit with somewhat irregular patterns) in *B. plebeius* abundance that corresponded to removal of seaweed from the diet, suggesting that its ability to maintain high abundance was tied to the presence of this resource (Figure 2C). Even in mice that do not receive seaweed, the non-native, human-derived, *B. plebeius* colonizes at low levels, but frequently falls below the limit of detection, suggesting that it has limited access to additional resources in the mouse gut to enable its persistence.

### High colonization with 1% resource advantage requires competition with endogenous commensals

To understand how *B. plebeius* can colonize abundantly in the mouse gut when seaweed is present as 1% (10 g/kg) of the diet, we make an argument using a mass balance. Assuming the mass of an individual cell of *B. plebeius* is 1 pg, obtaining the observed levels of *B. plebeius* in the mouse gut after seaweed treatment (approximately 10^10^ cells/g feces) requires a conversion efficiency on the order of 100% (10^10^ cells/g feces = 10^-2^ g cells/g feces = 1 g cells/g seaweed * 0.01 g seaweed/g feces). The concentration of seaweed by the time it reaches the colon is likely much greater than 1% (maybe 10% or more) as the host will absorb nutrients from starch (at 40%) and casein (at 20%) in the diet. However, achieving an efficiency of substrate usage between 10 and 100% is still surprising (see Supplemental Information). These observations together necessitate that *B. plebeius* is using alternative resources in the mouse gut *and* that it is more effective at accessing these resources in the presence than in the absence of seaweed. From a kinetic perspective, this scenario is easy to imagine: initially by increasing abundance on seaweed substrates, cells of *B. plebeius* have a numerical advantage in competing for other resources (see Supplementary Information).

This result also necessarily predicts that *B. plebeius* competitively inhibits the growth of other organisms if the total size of the ecosystem remains unchanged. The total abundance of bacteria in the system appears to remain unchanged in the presence or absence of seaweed as measured by qPCR. Keeping this in mind, as *B. plebeius* increases in abundance in the system, we expect many other sequence variants to decrease, and indeed, there is an enrichment for negative correlations with *B. plebeius* in the seaweed-treated mice compared to the untreated mice (309 negative correlations/328 total correlations with p < 0.01 for seaweed and pulsed groups and 9 negative correlations/30 total correlations with p < 0.01 for the *B. plebeius* only group, – see Methods for details). There is also a slight enrichment for negative correlations in the continuous seaweed group compared to the pulsed group.

There were few sequence variants that positively correlated with *B. plebeius* abundance. We hypothesized that the abundance of organisms with similar abundance and dynamic patterns might be constrained by the same limiting resource. We found that *A. muciniphila* significantly positively correlated with *B. plebeius* in two of the mice, one from the pulsed and one from the continuous seaweed diets (Figure 4A), and had no temporal relationship in the absence of seaweed. The remaining seaweed-treated mice both exhibited a decrease in *A. muciniphila* abundance over time, with no significant correlation to *B. plebeius*. Because *A. muciniphila* specializes in mucin degradation (Derrien, 2004), we inferred that both organisms use mucin for growth, indicating both a resource-based niche and interaction with the mucus layer. Intriguingly, increases in *A. muciniphila* abundance was associated with even greater increases in *B. plebeius* abundance (i.e. the slope is greater than 1) (Figure 4A), suggesting that *A. muciniphila* may in some way facilitate the growth of *B. plebeius.* Such growth facilitation has been shown previously for *A. muciniphila* grown with other gut commensals, including *B. vulgatus*, an organism incapable of using mucin on its own (Png et al., 2010).

### Long-term *B plebeius* persistence is reduced in mice constantly fed seaweed

By providing access to a limited amount of resource, we could explain the high level colonization of *B. plebeius* in the short term. We investigated whether this advantage would hold in the long term as well. We examined the presence of *B. plebeius* in mice two months after cessation of the initial seaweed-feeding period. We collected baseline samples and resumed 1% seaweed treatment in the groups that originally received seaweed. Surprisingly, although we observed blooms (to nearly 50% of the community) of *B. plebeius* on resumption of seaweed feeding in mice originally on the pulsed diet, there was no response of *B. plebeius* in the mice originally on the continuous diet, and the median relative abundance (1.6e-5) was just above the limit of detection for these animals (Figure 3A and Figure 3B). There was a single mouse in the pulsed seaweed group that lost *B. plebeius* entirely, but there was still a significant difference in the levels of *B. plebeius* between the GC and GP groups (p = 0.02, n = 5 GC mice, n = 5 GP mice, Mann Whitney U Test) (Figure 3A), suggesting a differential effect from the initial dietary regime on the long term responsiveness of *B. plebeius* to seaweed amendment.

**Figure 3.**
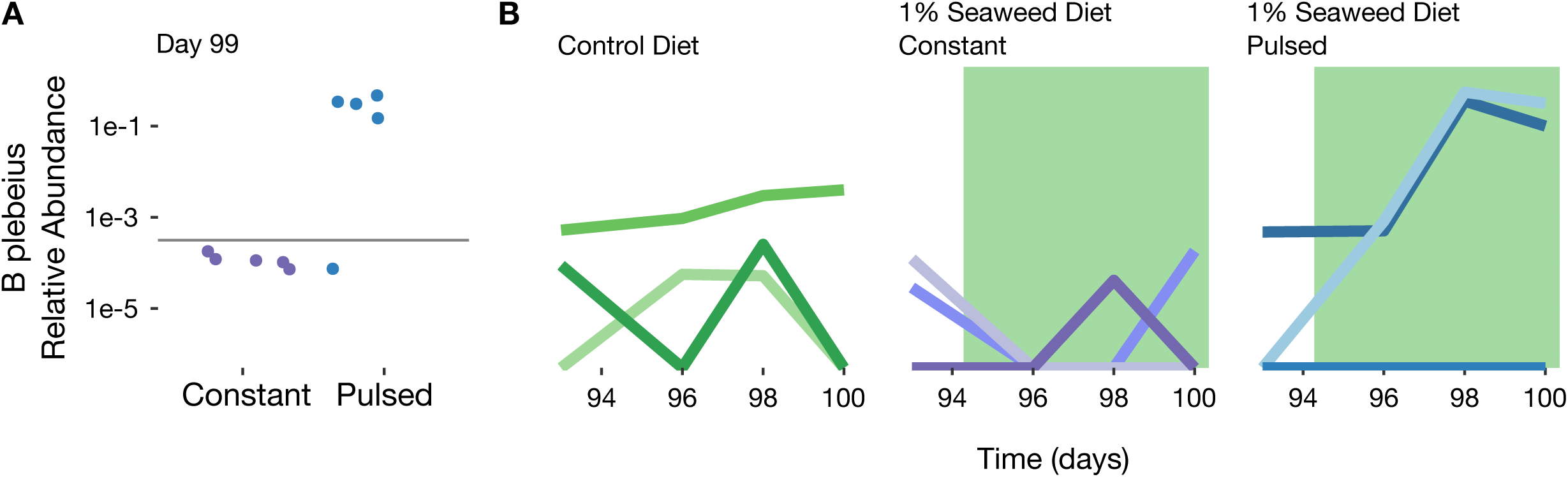
Long-term persistence of *B. plebeius* depends on initial diet regimen. (A) qPCR-based relative abundance of *B. plebeius* in constant and pulsed seaweed groups at indicated time point after 1% seaweed amendment. The lower limit of detection from non-specific amplification is marked with a gray line. (B) Relative abundance of *B. plebeius* over time in late time points on resumption of seaweed treatment after washout in *B. plebeius* only, constant, and pulsed seaweed groups.

### Diet-induced blooms of *B. plebeius* associate with increased IgA-binding

Recalling the apparent relationship of *B. plebeius* to *A. muciniphila*, and knowing *A. muciniphila* is heavily IgA-targeted in humans and mice (Palm et al., 2014), we wondered whether IgA-targeting, and thus immune activation against *B. plebeius* might be related to its disappearance from some of the mice, by inhibiting its ability to uptake seaweed polysaccharides through steric blocking or related mechanisms that prevent biofilm formation (Moor et al., 2017). Equally, IgA-targeting may be a mechanism for retaining bacteria while simultaneously controlling their growth (McLoughlin et al., 2016). High levels of IgA-binding do not necessarily imply that organisms are damaging, as introduction of highly IgA-coated species such as *A. muciniphila* and *C. scindens* can prevent enteropathy by colitogenic bacteria (Kau et al., 2015).

With these considerations in mind, we set out to identify whether *B. plebeius* was detected and targeted by IgA in the gut. Only in the case that *B.* plebeius generates a specific response by the immune system would we expect to find a signal of increased binding of *B. plebeius* relative to the rest of the microbiota, as polyreactive IgA tends not to preferentially bind *Bacteroidetes* (Bunker et al., 2017). Observing such a difference would provide a putative mechanism to attain control of *B. plebeius* abundance in the seaweed group at late time points, and potentially explain loss of *B. plebeius*.

To address whether the immune system may be involved in the apparent control of *B. plebeius*, we flow-sorted and sequenced bacteria that were highly and lowly IgA-bound, as described previously (Palm et al., 2014). At the late time points, *B. plebeius* was too low in abundance (< 1e-5) to detect in mice in the continuous seaweed treatment group. We focused our IgA-Seq efforts on late time point samples from mice originally on the pulsed and *B. plebeius* only groups, for which *B. plebeius* was detectable by qPCR. We failed to detect a signal of differential IgA-binding of *B. plebeius* in the *B. plebeius* only mice (Figure 4B), suggesting that IgA may not be important control mechanism for *B. plebeius* in these animals. However, in the pulsed mice that still had *B. plebeius*, there is a clear enrichment for this organism when it is present in the highly IgA-bound fraction (Figure 4B). In fact, it is more IgA-bound than any other member of the microbiota in one of these mice, suggesting that the binding was highly specific to this organism.

**Figure 4.**
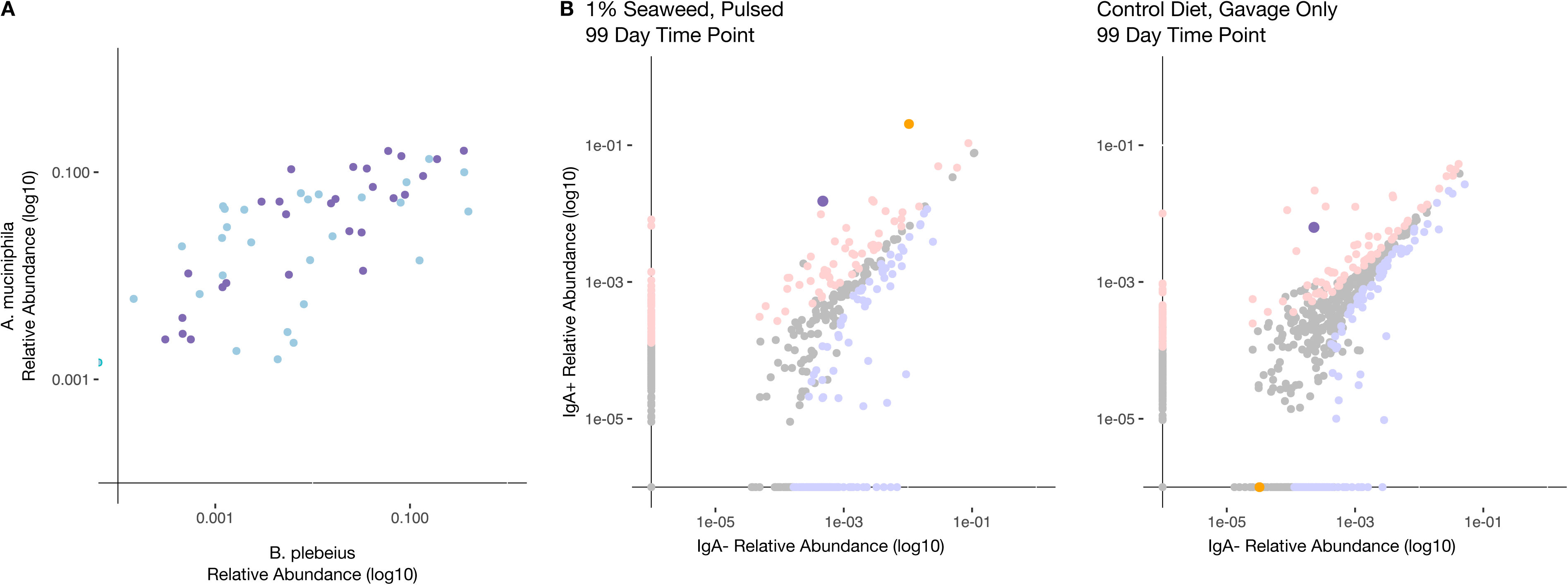
Endogenous competitors and the immune system may restrict long-term abundance. (A) Scatterplot of *B. plebeius* and *A. muciniphila* abundance over time for two animals having significant correlations between these bacteria (purple = an animal from the GC group, blue = an animal from the GP group). (B) Enrichment of sequence variants within the IgA+ and IgA-fraction; light red points are significantly enriched in the IgA+ fraction, light blue points are significantly enriched in the IgA-fraction. *B. plebeius* is indicated in red, *A. muciniphila* is indicated in green.

While we hesitate to generalize these observations, it seems possible that the IgA-binding of this organism is a response to its diet-induced abundance spike, as in general the abundance of this organism remains low in the absence of seaweed. The potential for diet-mediated IgA-targeting has previously been reported for *Enterobacteriaciae* (Kau et al., 2015). IgA-binding as a diet-microbe interaction for a gut commensal has important implications for diet-based manipulation of the microbiota, suggesting over-enrichment of certain organisms may trigger an immune response. Further, introducing small amounts of substrate to select for a specific microorganism may lead to its disproportionate expansion even in a highly competitive setting. These findings emphasize the importance of diverse substrates for maintaining a diverse microbiota.

## Discussion

Here, we provide a proof-of-concept of diet-based orthogonal niche engineering, whereby an organism is introduced with a tailored resource to establish an orthogonal niche in an intact community. Previous studies of the influence of diet on the gut microbiota have primarily focused on the alterations in the bacterial community and functional or metabolic shifts in response to dietary perturbations (Carmody et al., 2015; David et al., 2013, 2014; Desai et al., 2016; Turnbaugh et al., 2006, 2009). It has been documented that *Bacteroidetes* in particular display a hysteretic response (i.e. memory of past dietary exposures dampens future responses) to dietary changes (Carmody et al., 2015), which may arise as a consequence of an adaptive immune response to fluctuations in the abundance of these organisms. In all sampled mice that were originally on the seaweed diet, we observed boom-bust cycles of *B. plebeius*, particularly in the pulsed diet, suggesting a strong dependence on this resource when introduced into the system. We use a dynamical model to suggest that *B. plebeius* gains preferential access to mouse-gut endogenous resources in the presence of seaweed that lead to the prediction of increased competitive inhibition against native members of the gut microbiota. Temporal relationships between *B. plebeius* and *A. muciniphila* in mice treated with seaweed implied that *B. plebeius* might utilize mucin-derived substrates for growth. Use of mucin implies growth in the mucosa, which we expected would trigger immune responses against *B. plebeius*, particularly when it reaches high abundance. Important pathogens, including *C. difficile* and *S. enterica* have been shown to used mucin-derived substrates when metabolic networks among the native microbiota are disrupted, allowing them to expand to significant abundances within the community (Ng et al., 2013). This strategy may be widely used by commensal organisms, where disruption through dietary change promotes initial increases in abundance through preferential substrate access followed by increased competition for mucin glycoproteins.

In the 2-month follow-up time points, none of the mice originally on the constant seaweed diet again experienced blooms in *B. plebeius* following seaweed administration. We speculate that the consistently high levels of *B. plebeius* in the constant seaweed treatment group more potently triggers an immune response than in the mice not fed seaweed or those on the pulsed treatment, likely resulting in complete abolishment of its outgrowth at the later time points.

This diet-induced expansion of *B. plebeius* followed by eventual knockdown may apply more generally to gut commensal organisms. In the context of the dampening in response to dietary fluctuations observed previously (Carmody et al., 2015), the mechanism may be one in which diet-induced expansions of *Bacteroidetes* on one diet trigger an immune response that in subsequent exposures to the diet dampen the ability of these organisms to respond to substrate by coating them in antibodies. Extending this analysis to the generic case of changing diets, IgA-binding of commensals may be a strategy by which the host constrains outgrowth of strains able to grow on newly introduced dietary substrates (Kau et al., 2015). Although IgA-binding may disfavor expansion of a bacterial commensal, it may also enhance its retention in the system by providing a scaffold for adhesion (McLoughlin et al., 2016), which could be considered a feature for maintaining a commensal at non-disruptive levels within a host.

The heterogeneity in IgA-binding of strains across humans may be a consequence of dietary differences determining which organisms have outgrown historically within a person. Many PULs are carried on mobile elements, and so transfer frequently between strains of *Bacteroides* (Grondin et al., 2017; Jiang et al., 2017). The previously observed differences in antigenicity across strains of *B. fragilis* (Palm et al., 2014) suggests that this phenotype has a genetic basis, which, in contrast to classical virulence genes, might be rooted in the growth response of organisms to dietary substrate, a trait that can differ even between closely related strains because of the presence or absence of these PULs.

In general, dietary fiber seems to protect the mucus barrier from degradation by gut bacteria (Desai et al., 2016). However, seaweed as a substrate is unique: the sulfate groups on agar and porphyran, when cleaved, can be reduced to sulfides. Reduction of sulfates to sulfides by gut bacteria has previously been shown to deplete the mucus barrier (Ijssennagger et al., 2015), which may increase exposure of bacteria to patrolling immune cells, boosting the likelihood that the then-abundant organisms are targeted. However, fiber-induced mechanical stress to epithelial cells may in general be sufficient to disrupt the mucus layer (Miyake et al., 2006), such that the interaction between a particular dietary substrate and growth of organisms that thrive on this substrate will lead to preferential immune targeting of these organisms. The frequency-dependency of this response calls for more investigation.

These results provide a first proof of concept for orthogonal niche engineering as an approach to introducing bacteria into complex communities. Rational synbiotic design will benefit from considerations of the interaction of introduced prebiotics and probiotics with the rest of the biotic and abiotic community. In this system, there is very clear positive selection for growth on seaweed given that the genes were transferred from a marine bacterium to a gut commensal bacterium. But having these genes appears to be able to induce negative selection through IgA-targeting when the substrate is abundantly available. Inducing a robust and specific immune response may be a desirable feature of a microbial therapeutic, allowing for reversible engraftment. It will be important to consider how off-target effects through species-species, species-diet, and species-host interactions can alter these responses.

## Methods

### Animals

Six-week old female C57BL/6 wild type and outbred Swiss Webster mice (Taconic, Germantown, NY) were housed and handled in Association for Assessment and Accreditation of Laboratory Animal Care (AAALAC)-accredited facilities using techniques and diets including *Bacteroides plebeius* as specifically approved by Massachusetts Institute of Technology’s Committee on Animal Care (CAC) (MIT CAC protocol # 0912-090-15 and 0909-090-18). The MIT CAC (IACUC) specifically approved the studies as well as the housing and handling of these animals. Mice were euthanized using carbon dioxide at the end of the experiment.

### *B. plebeius* culture

*B. plebeius* DSM 17135 was obtained from the DSMZ and cultured as specified. Briefly, colonies were obtained by the streak method on PYG (Modified) Agar incubated at 37°C in a Coy Anaerobic Chamber (Grass Lake, MI) for up to 48 hours. After the appearance of colonies, single colonies were inoculated in PYG (Modified) medium and incubated overnight prior to gavage or nucleic acid extraction. 25% glycerol stocks of *B. plebeius* were made and stored at -80°C and streaked onto fresh medium for revival and colony picking.

### Seaweed Diet Experiment

C57BL/6 mice were used in the initial studies in Figure 1 in which 10 mice were fed with a custom chow diet (Bio-Serv, Flemington NJ) containing 1% raw seaweed nori (Izumi Brand) and 10 mice had a standard control diet (Product# F3156, AN-93G, Bio-Serv, Flemington NJ). Animals were co-housed for 6 days after arrival in the MIT animal facilities, and singly housed after separation into the seaweed treatment and control groups. Fresh fecal samples were obtained within an hour daily for all animals in all groups. Fecal samples were collected into anaerobic 25% glycerol containing 0.1% cysteine, and transferred immediately to dry ice before being stored at -80°C prior to nucleic acid extraction.

### *B. plebeius* Gavage Experiment

For the *B. plebeius* gavage experiments, Swiss Webster mice were co-housed for 6 days after arrival in MIT animal facilities, and 5 mice per group were co-housed by treatment on initiation of the experiment. Fecal samples were collected for each animal for three days prior to the initiation of the experimental protocol. At day 0, mice in the *B. plebeius* gavage groups were gavaged only once with approximately 10^7^ CFUs of *B. plebeius* DSM 17135 culture in 250 μl volumes of PYG (Modified) media, and control groups were gavaged only once with 250 μl sterile PYG (Modified) media. All groups treated with seaweed received a custom chow diet containing 1% seaweed nori at the initiation of experiments. All fecal samples were collected as described for the initial seaweed diet experiment.

### Nucleic Acid Extraction

DNA from fecal samples and bacterial cultures was extracted using the MoBio High Throughput (HTP) PowerSoil Isolation Kit (MoBio Laboratories, Inc., now Qiagen) with minor modifications. Briefly, samples were homogenized with bead-beating and then 50 μl Proteinase K (Qiagen) added and samples were incubated in a 65°C water bath for 10 minutes. Samples were then incubated at 95°C for 10 minutes to deactivate the protease. All other steps remained the same.

### 16S Library Preparation and Sequencing

Libraries for paired-end Illumina sequencing were constructed using a two-step 16S rRNA PCR amplicon approach as described previously with minor modifications (Preheim et al., 2013). In order to account for cross-sample and buffer contamination, triplicate negative controls comprising resistant fraction extraction blanks, nucleic acid extraction blanks, and PCR negatives were included during library preparation and samples were randomized across the plate. The first-step primers (PE16S_V4_U515_F, 5′ ACACG ACGCT CTTCC GATCT YRYRG TGCCA GCMGC CGCGG TAA-3′; PE16S_V4_E786_R, 5′-CGGCA TTCCT GCTGA ACCGC TCTTC CGATC TGGAC TACHV GGGTW TCTAA T 3′) contain primers U515F and E786R targeting the V4 region of the 16S rRNA gene, as described previously (Preheim et al., 2013). Additionally, a complexity region in the forward primer (5′-YRYR-3′) was added to help the image-processing software used to detect distinct clusters during Illumina next-generation sequencing. A second-step priming site is also present in both the forward (5′-ACACG ACGCT CTTCC GATCT-3′) and reverse (5′-CGGCA TTCCT GCTGA ACCGC TCTTC CGATC T-3′) first-step primers. The second-step primers incorporate the Illumina adapter sequences and a 9-bp barcode for library recognition (PE-III-PCR-F, 5′-AATGA TACGG CGACC ACCGA GATCT ACACT CTTTC CCTAC ACGAC GCTCT TCCGA TCT 3′; PE-III-PCR-001-096, 5′-CAAGC AGAAG ACGGC ATACG AGATN NNNNN NNNCG GTCTC GGCAT TCCTG CTGAA CCGCT CTTCC GATCT 3′, where N indicates the presence of a unique barcode.

Real-time qPCR before the first-step PCR was done to ensure uniform amplification and avoid overcycling all templates. Both real-time and first-step PCRs were done similarly to the manufacturer’s protocol for Phusion polymerase (New England BioLabs, Ipswich, MA). For qPCR, reactions were assembled into 20 μL reaction volumes containing the following: DNA-free H_2_O, 8.9 μL, HF buffer, 4 μL, dNTPs 0.4 μL, PE16S_V4_U515_F (3 μM), 2 μL, PE16S_V4_E786_R (3 μM) 2 μL, BSA (20 mg/mL), 0.5 μL, EvaGreen (20X), 1 μL, Phusion, 0.2 μL, and template DNA, 1 μL. Reactions were cycled for 40 cycles with the following conditions: 98**°** C for 2 min (initial denaturation), 40 cycles of 98 C for 30 s (denaturation), 52**°** C for 30 s (annealing), and 72**°** C for 30s (extension). Samples were diluted based on qPCR amplification to the level of the most dilute sample, and amplified to the maximum number of cycles needed for PCR amplification of the most dilute sample (18 cycles, maximally, with no more than 8 cycles of second step PCR). For first step PCR, reactions were scaled (EvaGreen dye excluded, water increased) and divided into three 25-μl replicate reactions during both first- and second-step cycling reactions and cleaned after the first-and second-step using Agencourt AMPure XP-PCR purification (Beckman Coulter, Brea, CA) according to manufacturer instructions. Second-step PCR contained the following: DNA-free H_2_O, 10.65 μL, HF buffer, 5 μL, dNTPs 0.5 μL, PE-III-PCR-F (3 μM), 3.3 μL, PE-III-PCR-XXX (3 μM) 3.3 μL, Phusion, 0.25 μL, and first-step PCR DNA, 2 μL. Reactions were cycled for 10 cycles with the following conditions: 98**°** C for 30 s (initial denaturation), 10 cycles of 98**°** C for 30 s (denaturation), 83**°** C for 30 s (annealing), and 72**°** C for 30s (extension). Following second-step clean-up, product quality was verified by DNA gel electrophoresis and sample DNA concentrations determined using Quant-iT PicoGreen dsDNA Assay Kit (Thermo Fisher Scientific). The libraries were multiplexed together and sequenced using the paired-end with 250-bp paired end reads approach on the MiSeq Illumina sequencing machine at the BioMicro Center (Massachusetts Institute of Technology, Cambridge, MA).

### 16S rDNA Sequence Data Processing and Quality Control

Paired-end reads were joined with PEAR (Zhang et al., 2014) using default settings. After read joining, the complexity region between the adapters and the primer along with the primer sequence and adapters were removed. Sequences were processed batchwise using the DADA2 (Callahan et al., 2016) pipeline in R, trimming sequences to 240 bp long after quality filtering (quality trim Q10) with maximum expected errors set to 1. A final sequence variant table combining all sequencing data was generated using DADA2. Sequence variants were classified using RDP (Maidak et al., 1996; Wang et al., 2007). The resulting count tables were used as input for analysis within R.

### qPCR

qPCR was carried out as described in the **16S rDNA Library Preparation and Sequencing** section. For quantification of *B. plebeius*, primers were designed to target the beta-porphyranase A gene (BACPLE_01693) in the porphyran degradation PUL: GH86 F: 5’-TCGAA TGTCA CAAAG CGTTC-3’ and GH86 R:

5’-ATGGA CGGGA CATTC TGTTC-3’. For direct quantification of *B. plebeius* abundance, a nucleic acid standard curve was prepared using 10-fold dilutions of nucleic acids extracted from *B. plebeius* overnight cultures quantified by NanoDrop after RNAse treatment. Mann Whitney U test was used to identify differences in *B. plebeius* abundance between treatment and control groups.

### IgA Sorting

Pre-weighed frozen fecal samples in glycerol were thawed at 4°C and then homogenized using a handheld homogenizer and pestles (Kimble Chase Kontes) at a final dilution in sterile PBS of 100 mg per ml. Samples were processed as described previously (Palm et al., 2014). After homogenization, samples were centrifuged at 50 × g at 4°C for 15 minutes, then washed three times in 1 ml PBS/1% BSA at 8000 × g for 5 minutes each. The pre-sort fraction was collected as 20 μL after resuspension prior to the final wash and stored at -80°C. The cell pellet was resuspended in 25 μL of 20% Normal Rat Serum (Jackson Immunoresearch) in PBS/1%BSA and incubated for 20 minutes on ice. After incubation, 25 μL 1:12.5 _-mouse-IgA-PE (EBioscience, clone mA-6E1) was added to each sample and samples incubated on ice for 30 minutes. Samples were washed three times in 1 ml PBS/1% BSA as above, and finally resuspended in PBS/1% BSA and transferred to blue filter cap tubes (VWR 21008-948) for flow sorting. An average of 500,000 cells from the IgA-positive and IgA-negative bacteria were sorted in triplicate into sterile microcentrifuge tubes on the BD FACSAria II at the MIT Koch Institute Flow Cytometry Core (Cambridge, MA). Samples were centrifuged, supernatant removed, and stored at -80°C until nucleic acid extraction using the DNeasy UltraClean Microbial Kit (Qiagen).

### 16S rDNA Data Analysis

A DADA2 sequence variant table including all experiments was imported into R and analyzed using phyloseq (McMurdie and Holmes, 2013) and custom R scripts.

Spearman correlations between *B. plebeius* and all other sequence variants in the time series were determined for each of the two mice across three groups. The number of sequence variants with negative or positive correlations and unadjusted p-values less than 0.01 were determined for each group. The Fisher Exact test was used to determine whether there were differences in the proportion of positive and negative correlations between the groups for each pairwise comparison of groups.

### IgA Data Analysis

We used DESeq2 to detect differences in abundance between IgA-bound and IgA-unbound fractions (McMurdie and Holmes, 2014). Using DESeq2 as applied to microbiome count data, OTUs that had an adjusted p-value of < 0.05 (Wald Test with Benjamini-Hochberg correction) between the IgA+ and IgA-fractions were considered significantly differentially abundant between the fractions.

### Model Construction and Analysis

See Supplementary Information for the chemostat model for bacterial growth. Code for model simulations were implemented in MATLAB R2017B.

